# enviLink: A database linking contaminant biotransformation rules to enzyme classes in support of functional association mining

**DOI:** 10.1101/2021.05.20.442588

**Authors:** Emanuel Schmid, Kathrin Fenner

## Abstract

**Motivation:** The ability to assess and engineer biotransformation of chemical contaminants present in the environment requires knowledge on which enzymes can catalyze specific contaminant biotransformation reactions. For the majority of over 100’000 chemicals in commerce such knowledge is not available. Enumeration of enzyme classes potentially catalyzing observed or *de novo* predicted contaminant biotransformation reactions can support research that aims at experimentally uncovering enzymes involved in contaminant biotransformation in complex natural microbial communities.

**Database:** enviLink is a new data module integrated into the enviPath database and contains 316 theoretically derived linkages between generalized biotransformation rules used for contaminant biotransformation prediction in enviPath and 3^rd^ level EC classes. Rule-EC linkages have been derived using two reaction databases, i.e., Eawag-BBD in enviPath, focused on contaminant biotransformation reactions, and KEGG. 32.6% of identified rule-EC linkages overlap between the two databases, whereas 40.2% and 27.2%, respectively, are originating from Eawag-BBD and KEGG only.

**Implementation and availability:** enviLink is encoded in RDF triples as part of the enviPath RDF database. enviPath is hosted on a public webserver (envipath.org) and all data is freely available for non-commercial use. enviLink can be searched online for individual transformation rules of interest (https://tinyurl.com/y63ath3k) and is also fully downloadable from the supporting materials (i.e., Jupyter notebook “enviLink” and tsv files provided through GitHub at https://github.com/emanuel-schmid/enviLink).

## Introduction

Refined understanding of contaminant degradation in environmental microbial communities depends on knowledge about catalyzing enzymes. For co-metabolic transformation at low substance concentrations that knowledge is hardly available. Available experimental approaches (gene knock outs or overexpression) are very costly and labor-intensive and therefore rely on strong hypotheses about potential enzyme candidates. To identify such potential enzyme candidates, functional association mining between metatranscriptomic or -genomic profiles and contaminant biotransformation information (i.e., rate constants and reaction pathway) has been suggested as a promising way forward ^1–3^. However, association mining suffers from low significance due to massive multiple hypothesis testing unless the range of enzymes plausibly catalyzing a given, observed transformation reaction can be restricted.

Currently available tools that, given a transformation reaction, allow predicting potentially catalyzing enzymes (or enzyme-encoding genes) are E-zyme/E-zyme2 ^4, 5^ and BridgIT ^6^. One obvious drawback for their application to contaminant biotransformation reactions is that both tools are trained on KEGG data only. KEGG very extensively covers reactions associated with primary metabolism and secondary metabolism of natural products, but only contains limited information on contaminants.

Eawag-BBD instead exclusively contains information on experimentally observed contaminant biotransformation reactions ^7^. These have served as a basis for deriving a set of manually curated generalized biotransformation rules (btrules) which are used for *de novo* contaminant pathway prediction ^8^. Most contaminant biotransformation reactions in Eawag-BBD are annotated with an EC number, which has been manually extracted by a data curator from the original publication reporting the experimental evidence. Most reactions are annotated with a 4^th^ or 3^rd^ level EC number (44.2% and 43.3%, respectively). The remaining 2^nd^ and 1^st^ level annotations are based on educated guesses of the data curators rather than actual experimentally proven linkages (personal communication, Prof. Lynda Ellis). Both, Eawag-BBD and Eawag-PPS have recently been implemented in a more flexible and state-of-the-art successor system called enviPath ^9^.

In developing enviLink, the database presented here, we therefore used reactions and their experimentally associated enzymes from both Eawag-BBD and, for completion, KEGG. We derived linkages between generalized biotransformation rules and 3^rd^-level EC classes rather than between actual reactions and 4^th^ level EC classes (as in BridgIT or E-zyme) for two reasons. First, given the enormous structural diversity of synthetic chemicals, the number of experimentally validated enzyme-reaction associations for contaminants is simply too low to derive a finer linkage scheme and validate it. Second, for the purpose of functional association mining, there is no need to target one enzyme only, but rather the goal is to produce a reasonably restricted, yet comprehensive list of suspect enzymes.

## Methods

The workflow for creating enviLink included three major steps (see Figure 1): (i) “in silico” reaction of Eawag-BBD and KEGG substrates against all Eawag-BBD biotransformation rules (btrules); (ii) Comparison of “in silico” generated reaction pairs (i.e., substrate(s) and product(s)) with Eawag-BBD or KEGG database reactions to find matching reactions; and (iii) generation of rule-enzyme links by extracting enzyme class of matching reactions and associating them with the btrule that predicted this reaction. Finally, to derive linkages between generalized biotransformation rules and 3^rd^-level EC classes, 4^th^-level EC numbers were summarized into the corresponding 3^rd^-level EC classes. All analyses were carried out separately for Eawag-BBD (1479 contaminant biotransformation reactions with 1301 associated EC classes) and KEGG (9952 reactions with 7007 associated EC classes, as of June 5^th^ 2020), and resulting links were compared as discussed below (note that for BBD 3^rd^ and 4^th^ level ECs were extracted, whereas for KEGG only the 4^th^ level ECs were considered). Details on each step of the workflow are given as Supporting Information in the form of interlinked Jupyter notebooks, which are available through GitHub (https://github.com/emanuel-schmid/enviLink). All data required to run the notebooks are available at this repository in the form of tsv files, but can alternatively also be downloaded following the code provided in the Jupyter notebooks.

**Figure 1:**
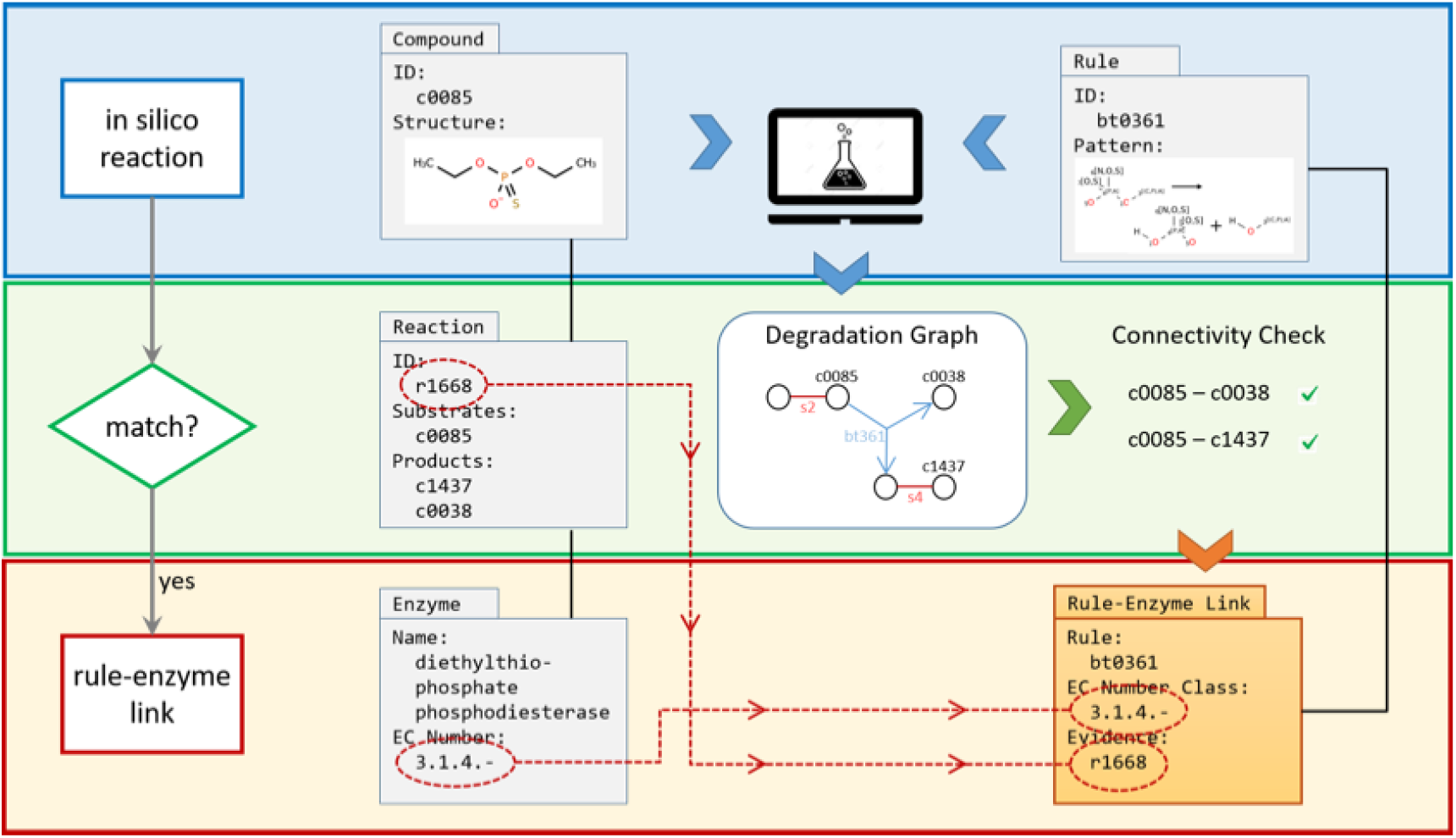
Overview of workflow to produce rule-enzyme linkages demonstrated for the example of Eawag-BBD and including three major steps: (i) Enumeration of “in silico” reactions by running all btrules against all BBD compound structures to produce predicted degradation graph for each BBD compound (blue upper panel, blue arrows), (ii) comparison of “in silico” prediction reactions from degradation graph with database reactions to check for matching reactions (green middle panel, green arrows), and (iii) generation of rule-enzyme links for matching reactions (red lower panel, orange arrow). Data entities are named in accordance with Eawag-BBD, and the example shown is taken from Eawag-BBD. In the degradation graph, blue lines stand for predicted reactions and red lines for standardizations. s2 and s4 represent standardizations and, in the specific case, stand for protonation/deprotonation reactions at differently substituted phosphate groups. Black connectors between data entities represent database relations, and red dashed connectors visualize the information flow from “Reaction”, “Enzyme” and “Rule” to yield entries for the new data entity “Rule-Enzyme Link” in enviLink.

## Results

Resulting linkages from Eawag-BBD and KEGG are accessible through the Jupyter notebook “enviLink results” at the GitHub repository. Alternatively, enviLink can be searched online at envipath.org for individual transformation rules of interest (see information given under “EC numbers” on the rule pages of the EAWAG-BBD package (https://tinyurl.com/y63ath3k)). Altogether 316 linkages between 169 btrules and 107 3^rd^ level EC classes were found and compiled in enviLink. For 39 btrules, no corresponding 3^rd^ level EC class could be identified. 32.6% of the identified rule-EC linkages overlap between the two databases, whereas 40.2% and 27.2%, respectively, are originating from either Eawag-BBD or KEGG only. The fact that more than one third of the linkages originate from Eawag-BBD exclusively demonstrates its unique information content with respect to contaminant biotransformation. One example of such an Eawag-BBD-exclusive linkage is the link between bt0241 and bt0242, two rules for hydroxylation of secondary and tertiary aliphatic groups, and 1.14.15, which contains monooxygenases using a reduced iron-sulfur protein as additional electron donor. Eawag-BBD contains literature entries reporting hydroxylating activity of camphor 5-monooxygenase (EC 1.14.15.1) on specific contaminants (e.g., adamantanone, tetralin) that are obviously not in the scope of KEGG and hence not reported therein.

In the “enviLink results” notebook, a histogram is provided showing how the linkages cover the space of btrules and 3^rd^ level EC classes. It can be observed that several 3^rd^ level EC are linked to multiple btrules (e.g., EC 1.14.12 is linked to bt0042, bt0072, bt0216 etc., which all encode for *vic*-dihydroxylation reactions at differently substituted aromatic rings). This illustrates that btrules in enviPath are divergent from the EC classification system in that they were optimized for specificity in contaminant biotransformation prediction^8, 10^.

Finally, to illustrate application of enviLink, consider the neonicotinoide acetamiprid, for which we observed enzymatic hydrolysis to the corresponding amide in activated sludge^11^. This reaction is predicted by bt0028 in enviPath, which in turn is linked to EC 4.2.1.-(hydrolases) in enviLink. When screening for associations between abundance of gene transcripts annotated to 4^th^ level EC classes belonging to 4.2.1.- and rate constants of acetamiprid biotransformation in activated sludge, nitrile hydratase transcript abundances (EC 4.2.1.84) showed significant correlations^1^. Indeed, own and literature evidence later confirmed that different nitrile hydratase homologs can turn over acetamiprid^1, 12^.

## Supporting information

Resulting linkages from Eawag-BBD and KEGG, tab delimited text

## Acknowledgements

We acknowledge financial support from the European Research Council under the European Union’s Seventh Framework Programme (ERC grant agreement 614768, PROduCTS).

